# Pollen tube growth and guidance: Occam’s razor sharpened on a molecular AGP Rosetta Stone

**DOI:** 10.1101/167890

**Authors:** Derek T. A. Lamport, Li Tan, Michael Held, Marcia J. Kieliszewski

## Abstract

Occam’s Razor suggests a new model of pollen tube tip growth based on a novel Hechtian oscillator that integrates: (1) a periplasmic AGP-Ca^2+^ calcium capacitor with tip-localised arabinogalactan glycoproteins (AGPs); (2) tip-focussed cytosolic Ca^2+^oscillations; (3) Hechtian strands evidence of adhesion between the plasma membrane and the cell wall of the growing tip. Thus Hechtian adhesion, as a piconewton force transducer, couples the internal stress of a rapidly growing wall to the plasma membrane. Such Hechtian transduction via stretch-activated Ca^2+^ channels and H^+^-ATPase proton efflux dissociating periplasmic AGP-Ca^2+^, creates a Ca^2+^ influx that activates exocytosis of wall precursors. In effect a highly simplified primary cell wall regulates its own synthesis and a Hechtian growth oscillator regulates overall tip growth. By analogy with the Rosetta Stone that translates trilingual inscriptions as a single identical proclamation, the Hechtian Hypothesis translates classical AGPs and their roles as a Ca^2+^ capacitor, pollen tube guide and wall plasticiser into a simple but widely applicable model of tip growth. Even wider ramifications of the Hechtian oscillator may implicate AGPs in osmosensing or gravisensing and other tropisms, leading us yet further towards the Holy Grail of plant growth.

## Introduction

In 1682 Nehemiah Grew in “The Anatomy of Plants” (1682) described stamens as male organs and their pollen as necessary for fruit production. Somewhat later Amici (1824) observed pollen germinating on the stigma and suggested that the pollen tube carried sperm cells to the ovule. Over fifty years ago (Mascarenhas & Machlis, 1962) a chemotropic dependence on Ca^2+^ for pollen tube growth and guidance became evident. Such growth is of particular interest as the pollen tube tip has the simplest primary cell wall consisting largely of highly methyl esterified pectic polymers and shows the fastest known tip growth rate that is generally not continuous but pulsatile (Pierson *et al*., 1995) or oscillatory with associated ion fluxes, notably H^+^ and Ca^2+^ (Feijo *et al*., 1995) of similar periodicity. However these ion fluxes are not in phase with tip growth rates (Michard *et al*., 2009) so causal relationships are not obvious (Messerli & Robinson, 2003; Holdaway-Clarke *et al*., 1997). With Occam’s Razor as a guide we view tip growth as a biological oscillator that depends on two novel components, namely classical AGPs and Hechtian adhesion sites. Based on the pH-dependent reversible binding of Ca^2+^ by AGPs, the Hechtian oscillator accounts for pollen tube H^+^ and Ca^2+^ ion currents as follows: H^+^ dissociates periplasmic AGP-Ca^2+^ thus increasing free cytosolic Ca^2+^ that coordinates exocytosis of cell wall precursors. However, cytosolic Ca^2+^ also depends on opening stretch-sensitive Ca^2+^ channels by tension from the growing wall transmitted to the plasma membrane by Hechtian adhesion sites. Because the new model includes two components, notably classical AGPs and Hechtian adhesion sites (Fig. 1) not previously considered as essential to models of extension growth, we suggest that overall tip growth consisting of the above components is a Hechtian oscillator. This model (Fig. 2) may also illuminate the vexed problem of cell extension dating back to Heyn’s identification of cell wall plasticity and via its hormonal regulation, as the primary factor in controlling growth by cell extension.

**Fig. 1.**
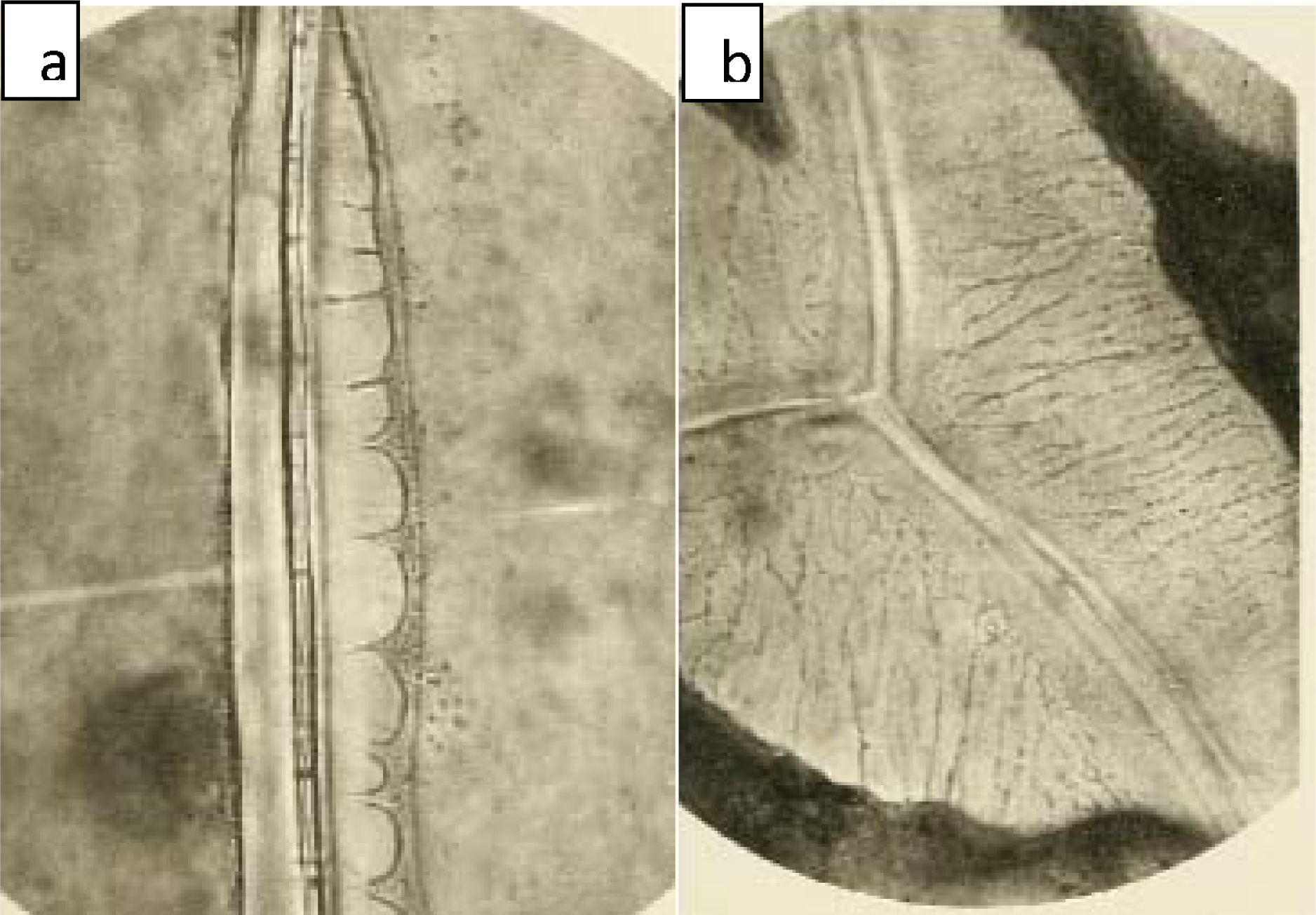
Hechtian strands of plasmolysed onion epidermis. These plasmolysed cells depict the founder event. Reprinted from (Hecht, 1912). A. Incipient plasmolysis of onion epidermal cells in 3% KNO_3_ B. Plasmolysis in 7% KNO_3_

AGPs, generally regarded as mysterious molecules (Pickard, 2013; Pereira *et al*., 2015) and ‘minor’ cell wall components are associated with many aspects of cell signalling (Seifert & Roberts, 2007) and cell expansion (Ding & Zhu, 1997). However, classical AGPs per se have not been viewed as essential regulatory components of pollen tube growth. Here we propose that classical AGPs based on their molecular properties and periplasmic location (Lamport et al. 2006), make a threefold contribution: Firstly as a primary source of cytosolic Ca^2+^ waves, secondly, as a pectic plasticizer and thirdly, as Ca^2+^ signposts to the ovule. These three inscriptions on an allegorical AGP molecular Rosetta Stone translate into general plant growth that depends on the remarkable chemical properties of AGPs summarised in Table 1 of (Lamport et al. 2014) and briefly as follows:

(1) AGP amino acid composition, rich in hydroxyproline (Lamport, 1970); (2) AGP O-Hyp glycosylation by small acidic arabinogalactan polysaccharides (Lamport, 1977) whose heterogeneity is generally exaggerated; (3) specific binding of Ca^2+^ by these glycomodules (Lamport & Varnai, 2013) amounting to a typical AGP Ca^2+^ content of ~1% w/w; (4) AGP location initially attached to the plasma membrane by a GPI anchor (Oxley & Bacic, 1999), but cleaved to allow incorporation into the growing wall; and finally (5) AGP molecular size of ~100kDa hence extrusion rather than simple diffusion of AGPs through the wall matrix after GPI-anchor cleavage (Lamport *et al*., 2006).

Several recent mathematical models predict oscillations in tip growth based on Ca^2+^-dependent vesicle recycling and tip plasticity (Zerzour *et al*., 2009; Hill *et al*., 2012). However, none include AGPs and Hechtian feedback as essential components. Our new model supplements these biophysical approaches with the evidence for a central role of AGPs in a biochemical model of tip growth as follows.

## A Hechtian model of pollen tube tip growth

Briefly, the new model (Fig. 2) proposes that a viscoplastic pectic cell wall mechanically coupled to the plasma membrane by Hechtian adhesion (Hecht, 1912) transmits wall strain to the PM and thus regulates H^+^, Ca^2+^ and other ion fluxes that regulate the exocytosis of wall precursors. Together they constitute a Hechtian oscillator (Fig. 2) consistent with Pickard’s pioneering work on mechanotransduction (Pont-Lezica *et al*., 1993). This explains how the rapidly growing cell wall of the pollen tube tip can act as a decision maker that regulates its own growth by Hechtian feedback. Such dramatic oscillatory growth (0.1-0.5 μm s^-1^) (Hepler *et al*., 2013) associated with active ion fluxes predominantly Ca^2+^ (Miller *et al*., 1992) H^+^, and K^+^ reflect Peter Mitchell’s chemiosmotic paradox (Mitchell, 1961): “ *Not only can metabolism be the cause of transport, but also transport can be the cause of metabolism”*. While there has been much effort to relate these ion fluxes to extension growth the Hechtian oscillator resolves the chemiosmotic paradox of the pollen tube by assigning specific roles to the Ca^2+^ and H^+^ ion currents essential for oscillations in tip growth (Hepler *et al*., 2006) driven by turgor pressure: Thus an initial acceleration of tip extension followed by exocytosis of wall precursors leads to deceleration, summarised in a simplified model (Fig. 2) partly based on Figure 3 of (Holdaway-Clarke & Hepler, 2003).

**Fig. 2.**
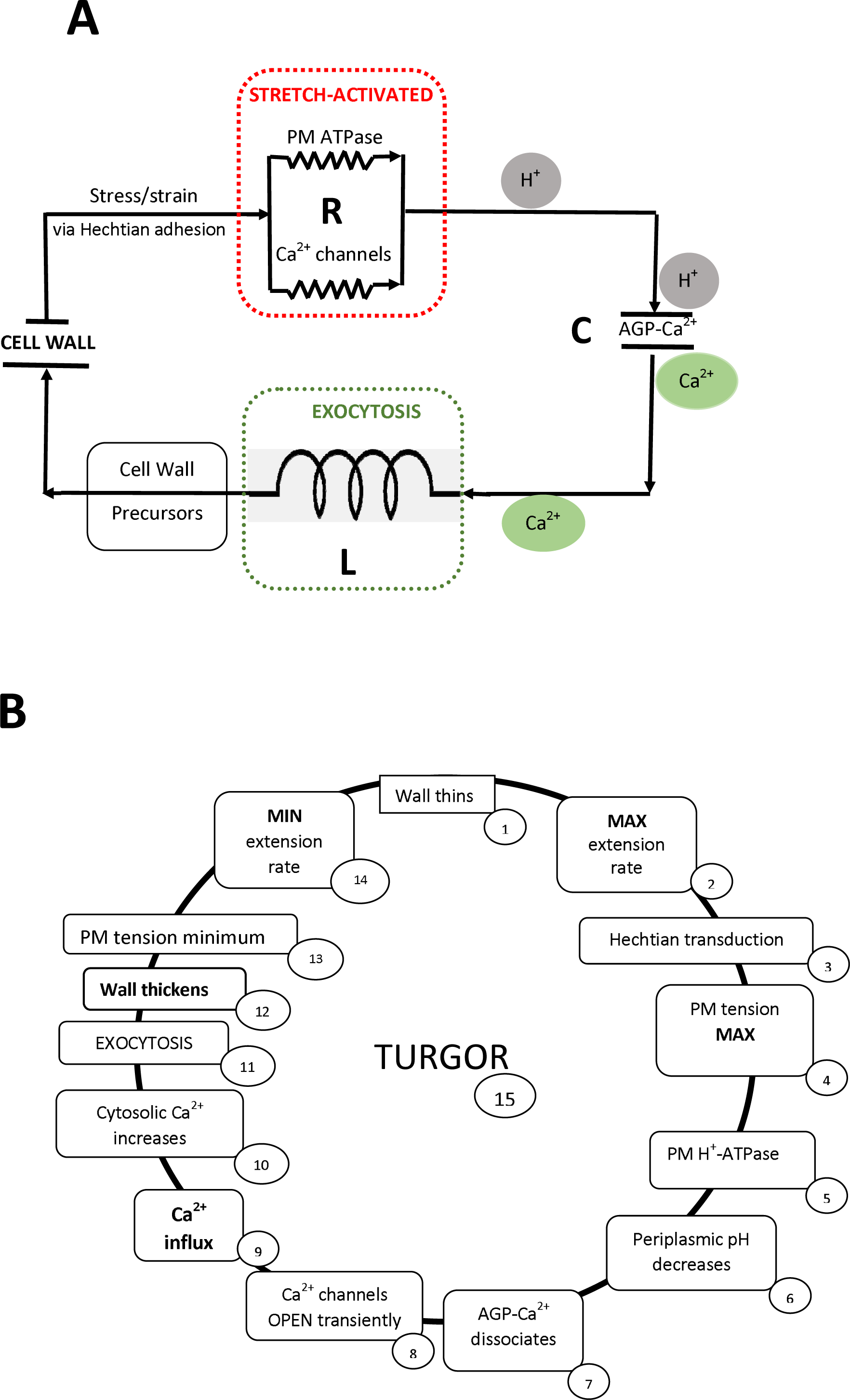
The pollen tube as a Hechtian <u>oscillator</u>. **A**. The Hechtian oscillator commemorates the contributions of Karl Hecht. Nominally the three simple terms R, L and C form the pollen tube tip growth oscillator that shows how the wall regulates its own growth by coupling H^+^ and Ca^2+^ ion currents: Turgor pressure charges the battery i.e. stretches the wall that transmits the resulting internal stress via Hechtian adhesion to the plasma membrane stretch-activated H^+^-ATPase and Ca^2+^-channels (resistance R). Together they work in parallel where H^+^ releases bound Ca^2+^ (capacitance C) to the Ca^2+^ channels hence a major source of cytosolic Ca^2+^ that activates exocytosis (inductance L) of wall precursors. Arguably these largely determine the frequency of tip growth oscillations; both auxin (Zerzour *et al*., 2009) and fusicoccin (Fricker *et al*., 1997) enhance oscillatory growth consistent with their activation of plasma membrane H^+^ ATPase. Hechtian strands are prominent features of many cells on plasmolysis including root hairs, pollen tubes, stomatal guard cells and green algae like Closterium (Domozych *et al*., 2003). This implies a global role for Hechtian adhesion as a stress-strain gauge that can also act as a sensor and regulator of turgor pressure. **B**. illustrates a single cycle of the oscillator based essentially on (Chebli & Geitmann, 2007; Hepler *et al*., 2013) arbitrarily divided into 14 stages beginning conveniently with rapid tip growth and transduction of wall stress. Although depicted as a simple cycle, most stages comprise critical control points with multiple inputs. Some stages remain to be defined such as cleavage of GPI-anchored AGPs via PLC activity (Dowd *et al*., 2006) that is possibly also stretch activated. Others such as PM ATPase activity (Certal *et al*., 2008) at the tip itself are surmised although activity is strongest just behind the tip at the pollen tube shank (Hepler *et al*., 2006). While ion fluxes are not precisely in phase with growth their synchronicity suggests a close relationship as inferred here:

1. Wall thins
2. Tube extension rate is inversely proportional to wall thickness(Kroeger *et al*., 2008)
3. Hechtian transduction transmits wall stress (Pont-Lezica *et al*., 1993)
4. Increases tension in plasma membrane tethered to cell wall: (Hecht, 1912).
5. Proton efflux via PM H^+^ATPase (Certal *et al*., 2008)
6. Decreases periplasmic pH
7. Dissociates periplasmic AGP-Ca^2+^ (Lamport *et al*., 2014; Lamport & Varnai, 2013)
8. Stretch-activated Ca^2+^ channels open(Ding & Pickard, 1993; Feijo *et al*., 1995; Dutta & Robinson, 2004)
9. Ca^2+^ influx via open Ca^2+^ channels
10. Cytosolic Ca^2+^ increases (Miller *et al*., 1992)
11. Activates exocytosis of wall precursors. (Camacho & Malho, 2003)
12. Wall thickness at tip increases (Picton & Steer, 1983; McKenna *et al*., 2009)
13. PM tension decreases
14. Minimum rate of tip extension (McKenna *et al*., 2009) Return to step 1.
15. Turgor drives tip extension (Hill *et al*., 2012) Note: Cytosolic Ca^2+^ recycles via Ca^2+^-ATPase mediated efflux (Schiott *et al*., 2004; Frey *et al*., 2015) or sequestration by golgi exocytotic vesicles.

### Proton flux

Ion fluxes drive all growth (Armstrong, 2015): Protons lead the way as the source of the chemiosmotic proton motive force (Mitchell, 1961) that generates ATP via mitochondrial F_l_F_0_ ATP synthase (Allegretti *et al*., 2015). However, in reverse the synthase pumps protons (Mazhab-Jafari *et al*., 2016). Thus plasma membrane H^+^ ATPase proton efflux is essential for maintaining the membrane potential and ion transport (Hepler *et al*., 2013). However, in growing pollen tubes unevenly distributed membrane H^+^-ATPases (Certal *et al*., 2008) result in a pronounced proton efflux at the tube shank with an apparent much smaller oscillatory influx at the tip (Feijo *et al*., 1999). A large proton efflux may explain the striking difference between the optimum extracellular pH of animal and plant cells (pH 7.4 and pH 5.5); this reflects the differing compositions of their extracellular matrix and their dynamic Ca^2+^storage that is largely peripheral in plant cells but mainly intracellular in animals. The lower external pH of plants reflects the low pK_a_ of uronic acids enabling pH-dependent uptake and release of Ca^2+^ from AGPs as a result of H^+^ ATPase activity. That also accounts for the massive proton efflux at the tube shank and ensures Ca^2+^ release from the abundant AGPs of stigmatic tissues hence a possible cooperative effect on the growth of other pollen tubes. (cf. (Lord, 2003)). The role of the less marked tip H^+^ influx (Feijo *et al*., 1999) is less clear as direct measurement cannot detect a much smaller H^+^ efflux into the nanometre dimensional domain of periplasmic AGP-Ca^2+^.

### AGP-Ca^2+^ as a primary source of cytosolic Ca^2+^

According to the prevailing view that ignores the largely methyl esterified status of tip pectin “ *Wall binding of Ca^2+^ accounts for the extracellular influx*” (Hepler *et al*., 2013) via open Ca^2+^ channels. Here we correlate the tip-focussed cytosolic Ca^2+^ oscillations with the presence of tip-localised AGPs (Figure 3A in (Mollet *et al*., 2002)) based on their pH-dependent dissociation (Lamport & Varnai, 2013) by plasma membrane H^+^-ATPase (Koji *et al*., 2012). The nanometre dimensions of periplasmic AGP-Ca^2+^ and its proximity to the proton source results in a significantly lower pH than the pH in muro due to rapid dissipation of the proton concentration by diffusion, dilution and buffering.

The advantage of AGP-Ca^2+^ as a source of cytosolic Ca^2+^ at the tip arises not only from its cell surface location a prime area of signal perception, but also from the ***paired glucuronic acid sidechains*** of AGP glycomodules (Fig. 3); these increase the total Ca^2+^ binding capacity of AGPs compared with the lower binding capacity of highly methyl esterified pectin that has largely unpaired galacturonic acid residues. Because the AGP glucuronic acid pK_a_ is lower than that of pectin galacturonic acid (Lamport & Varnai, 2013) AGPs bind Ca^2+^ more strongly at low pH. Thus periplasmic AGPs can also act as a sink for less firmly bound hence more easily released Ca^2+^ from pectin in the tip wall. Finally, the location of abundant periplasmic AGPs **confers a large kinetic advantage to an AGP-Ca^2+^** capacitor (Lamport & Varnai, 2013) that can readily supply the stretch-activated Ca^2+^ channels regulated by membrane tension (Dutta & Robinson, 2004) as follows:

**Fig. 3.**
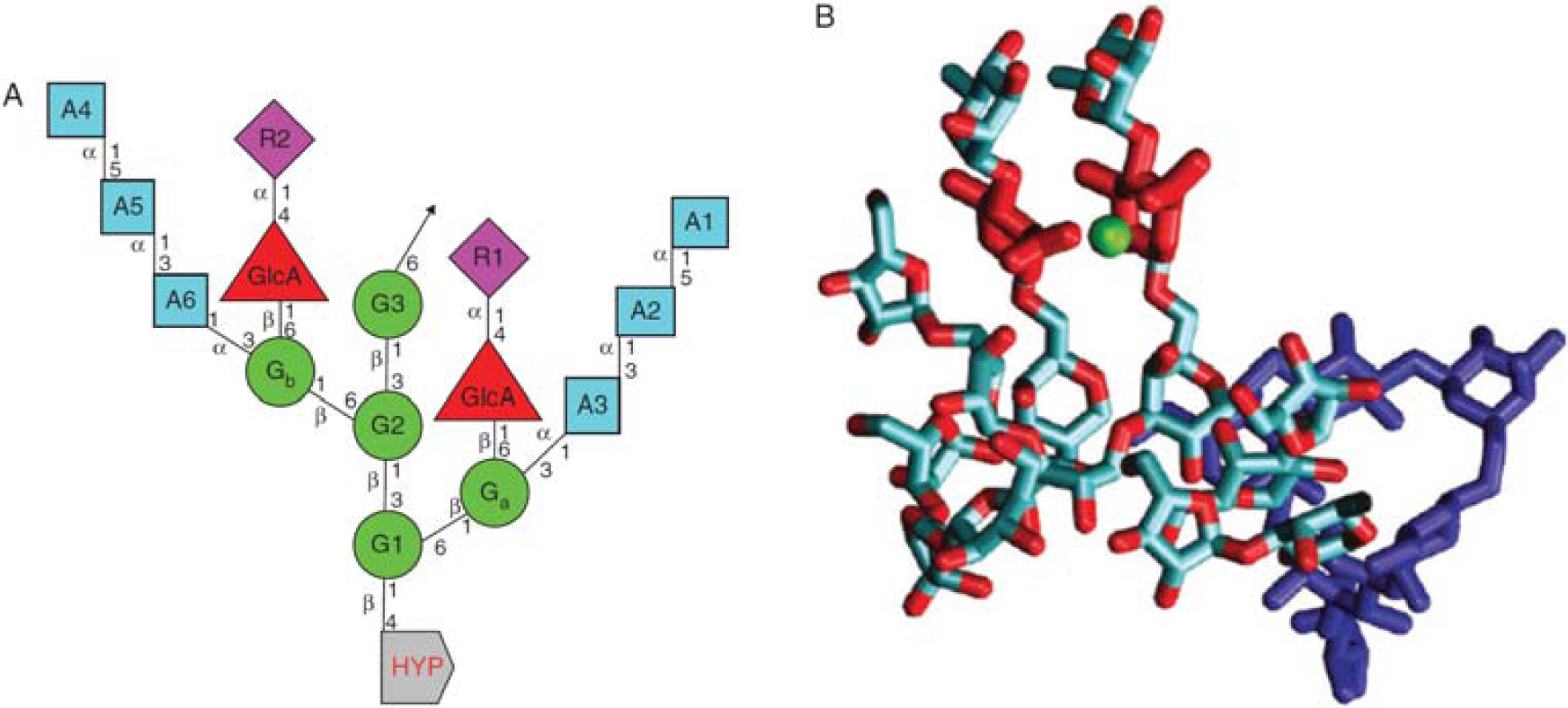
Repetitive AGP glycomodule. The 15-residue glycomodule structure that binds Ca^2+^ as depicted above corresponds to known highly expressed AGPs (Tan *et al*., 2010). (A) The linkage connectivity of sugars involved in the repetitive 15-residue consensus Hyp–arabinogalactan, a conserved structure based on a β-linked galactosyl trisaccharide with paired sidechains that bind Ca^2+^. A: 1–6 arabinose residues; G: 1–3 galactose mainchain residues; Ga and Gb: galactose sidechain residues; R1 and R2: rhamnose sidechain residues; GlcA: glucuronic acid sidechain residues; Hyp: hydroxyproline. (B) Three-dimensional molecular model simulating a Hyp–AG with bound Ca^2+^ Hyp–AG interferon Hyp-polysaccharide-1 (IFNHP1) with Ca^2+^ ions (green) bound by two glucuronic acid (GlcA) sidechains (red); the galactan backbone is in dark blue and sidechains in light blue. [Reprinted from (Lamport & Varnai, 2013)] See Supplementary Movie S1 in (Lamport & Varnai, 2013). Molecular dynamics simulation of Hyp-AG IFNHP1 (Gal backbone in blue) showing the conformational change on Ca^2+^ (green) binding by two GlcAs (red) over 500 ns. nphl2005-sup-0003-LegendMovieS1.doc

### Hechtian transduction

The significance of Hechtian adhesion evidenced by thread-like elastic extensions of the plasma membrane physically connecting the membrane to the cell wall of plasmolysed cells has remained obscure for more than a hundred years. (Fig. 1.) (Hecht, 1912). Hechtian adhesion is particularly evident in rapidly growing cell suspension cultures (Lamport, 1963) and during tip growth of root hairs (Volgger *et al*., 2010) and pollen tubes (Fig. 4.) (Parton *et al*., 2001). However absence of Hechtian adhesion from the pollen tube shank (Lord, 2003) confirms a significant role during tip growth further emphasised by its presence during tip growth even in chlorophycean algae like Closterium (Domozych *et al*., 2003). Such evolutionary conservation also supports a fundamental biological role of Hechtian adhesion in regulating plant growth inferred here: Stable Hechtian adhesion arises from strong molecular anchoring forces, most likely of AGPs and formins. Arguably AGP GPI lipids with an adhesion force of ~350 piconewtons (Cross et al. 2005), supplemented by formin transmembrane domains, enable the growing wall to transmit its stress/strain status at very low piconewton levels (cf. Buer et al. 2000) to the protoplast via multiple Ca^2+^ channels and H+-ATPases of the plasma membrane. High sensitivity Hechtian stress transducers are thus consistent with much evidence of stretch-activated Ca^2+^ channels (Dutta & Robinson, 2004) and a Hechtian “stress focussing” structure involving AGPs suggested earlier (Gens *et al*., 2000). Hypothetical wall-plasma membrane wall linkers proposed earlier (Pont-Lezica *et al*., 1993) involve specific candidates we identify here as AGP57C (At3g45230) and formin1 AtFH1; their well-defined molecular domains interact specifically with both plasma membrane and cell wall: AGP57C, an arabinoxylan-pectin-AGP glycoconjugate (APAP1) (Tan *et al*., 2013) has a C-terminal sequence directing GPI-anchor addition hence attachment to the plasma membrane, while the terminal rhamnose of an AGP glycomodule is attached to the reducing end of RGI a wall pectic polysaccharide. Formin1 has an N-terminal signal peptide followed by a transmembrane domain and a large extracellular domain anchored to the wall (Martiniere *et al*., 2011) most likely involving AGP glycomodules encoded by the Hyp glycosylation motif SPSALSPS.

**Fig. 4.**
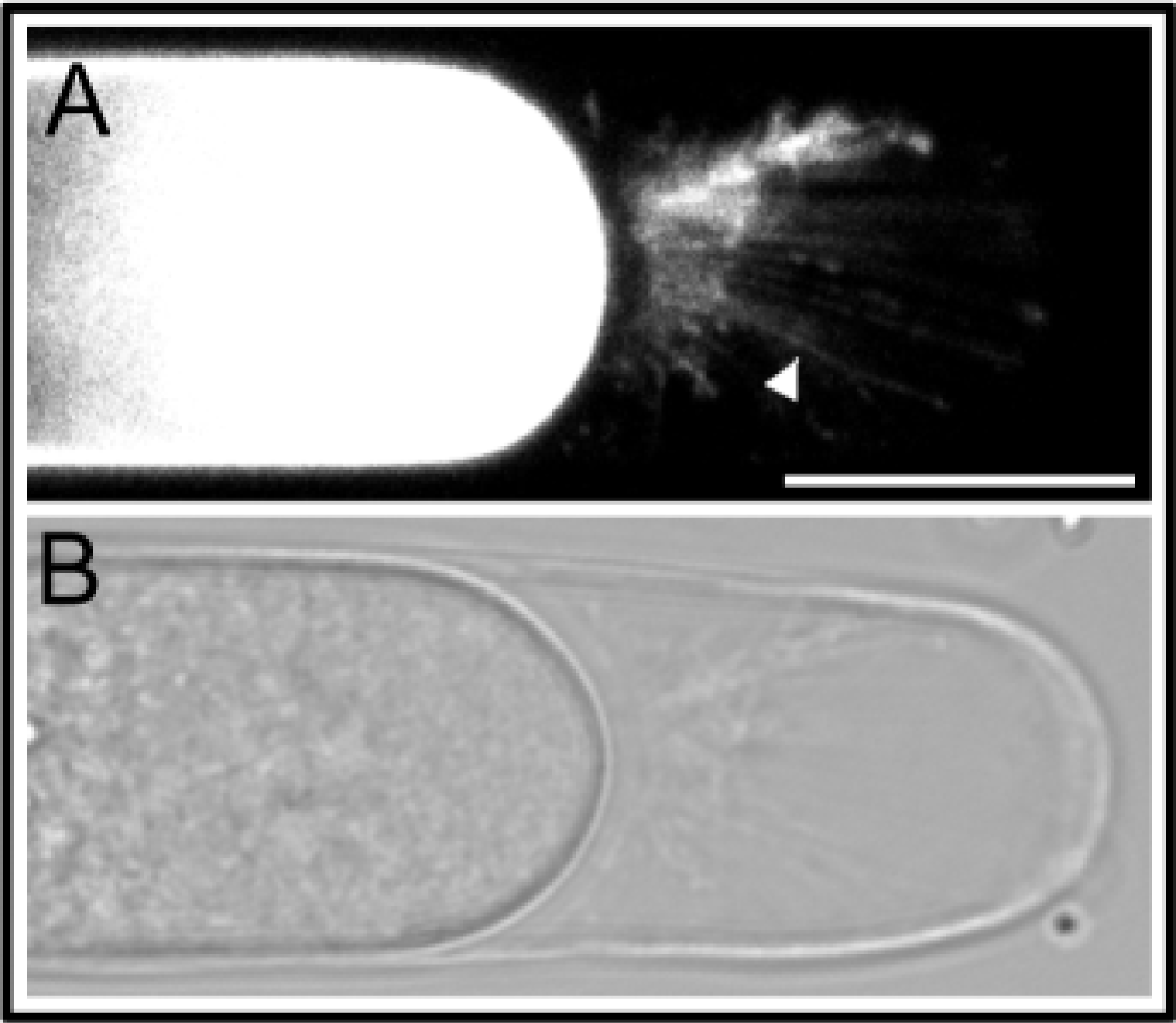
Plasmolysed pollen tube tip showing Hechtian strands. Plasmolysis of a *Lilium longiflorum* pollen tube loaded with the fluorescent dye FM4-64 five minutes after transfer to medium containing 0.3 M sorbitol and 2 μM FM4-64. (A) arrow head indicates fluorescent Hechtian strands. (B) Brightfield image. Bar = 15 μm. With permission from (Parton *et al*., 2001).

AGP57C and formin1 fulfil the criteria for bona fide crosslinks between a wall polysaccharide and the plasma membrane; at last providing tangible molecular evidence for Hechtian adhesion and its pivotal role in Hechtian signal transduction.

### Exocytosis of wall precursors

Exocytosis, the final stage of the secretory pathway from Golgi to cell surface, involves actin guidance and transport of exocytotic Golgi vesicles, docking and fusing with the plasma membrane. Although treated here as a single “component” of an oscillator, exocytosis involves many proteins and elevated levels of cytosolic Ca^2+^ at sites of pronounced exocytosis in growing pollen tubes (Camacho & Malho, 2003). Alteration of the Ca^2+^ gradient alters the pattern of exocytosis (Ge *et al*., 2007) suggesting the role of cytosolic Ca^2+^ as a coordinator of exocytosis consistent with the numerous Ca^2+^-dependent membrane processes (Luckey, 2008) and greatly decreased exocytosis when the Ca^2+^ chelator chlortetracycline decreased cytosolic Ca^2+^ (Reiss & Herth, 1978). While Ca^2+^-regulated exocytosis is a prime candidate for the regulation of oscillatory pollen tube growth, paradoxically some have concluded from the apparent lack of correlation between Ca^2+^ and secretion that although exocytosis may regulate oscillatory pollen tube growth, intracellular Ca^2+^ does not regulate oscillatory exocytosis (McKenna *et al*., 2009). Exocytosis, as a component of a Hechtian oscillator, connects H^+^ and Ca^2+^ ion gradients with cell wall tip growth (metabolism) and is thus a classic example of the Mitchell Paradox. Other models of tip growth based on ROP GTPases for example (Yan *et al*., 2009) did not include the cell wall or AGPs. However, due to technical difficulties the biochemical properties of the tip cell wall have received relatively little attention even though it is a major component of growth and its raison d’être.

### Cell wall plasticity

At the tip of the Lily pollen tube maximum thickness of the wall coincides with a significantly decreased rate of tip growth (Figure 4c in (McKenna *et al*., 2009)) which ***precedes*** a more rapid expansion (McKenna *et al*., 2009). As the wall thins its plastic extensibility increases with concomitant acceleration of the tip growth rate; further exocytosis restores wall thickness and thus decreases a growth rate that is inversely proportional to wall thickness (Kroeger *et al*., 2008). Evidently, wall plasticity plays a crucial role (Heyn, 1940) in determining rheology of the pollen tube tip primary cell wall. An explanation for the plasticity of primary walls in general is complicated by their wide range in composition with differing proportions of major components that include, cellulose, pectin, xyloglucan and structural protein (Fig. 5). However, the pollen tube tip is an ideal model of extension growth because it represents a simplified primary cell wall (originally defined by (Kerr & Bailey, 1934)) stripped down to a “single” major macromolecular pectin component. “Pectin is not just jelly” (Roberts, 1990) but forms a highly ordered composite of three major pectic polysaccharide domains, highly methylesterified polygalacturonide, RG-I, a highly methylesterified linear polygalacturonate and RG-II a complex rhamnogalacturonan with intermolecular borate crosslinked sidechains, unstable at low pH (Yapo, 2011). AGP α-L-arabinosyl sidechains may interact with pectin rhamnogalacturonan RG-II by competing with the terminal α-L-Gal of RG-II sidechain-A (Yapo, 2011) with concomitant disruption of its apiosyl borate ester intermolecular crosslink.

**Fig. 5.**
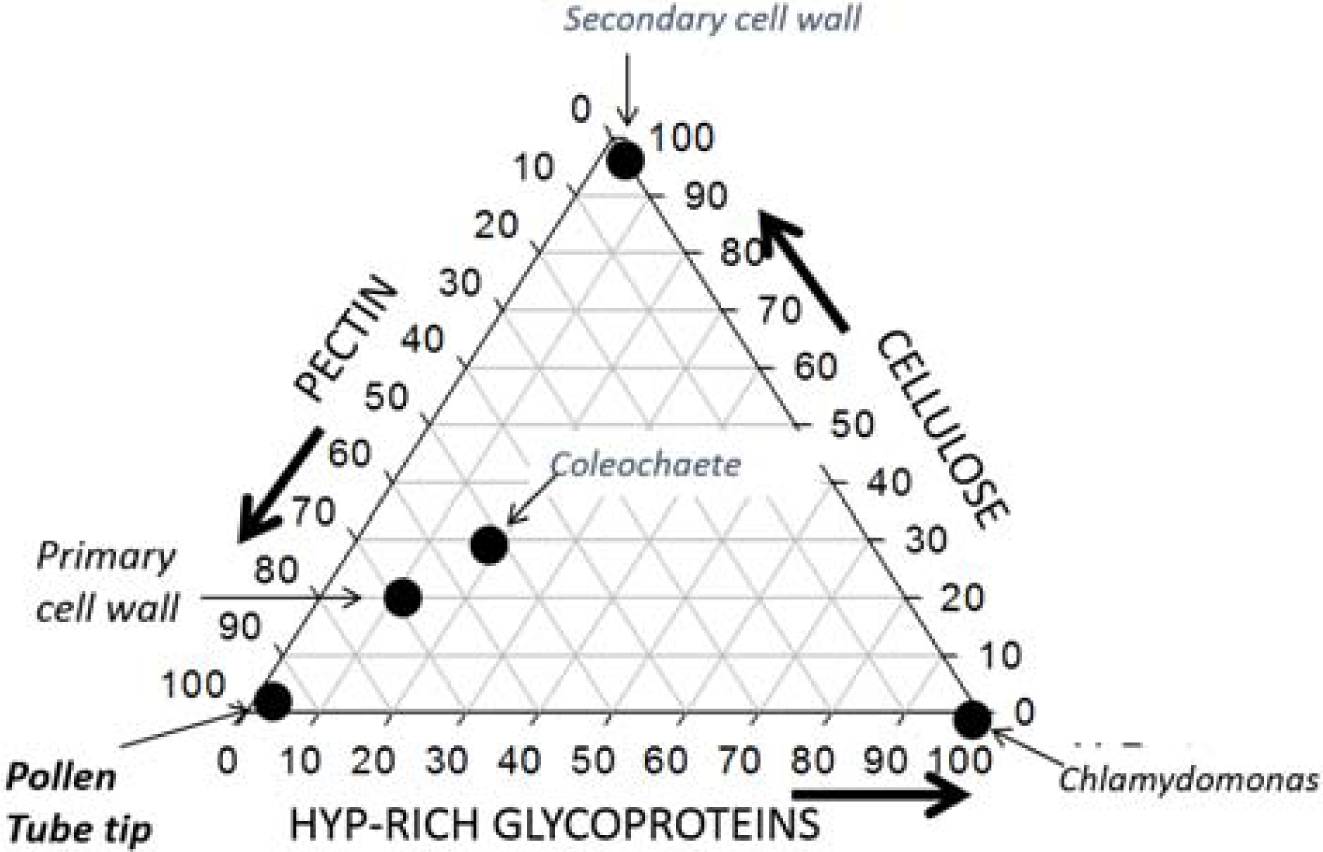
Comparison of cell wall compositions. Ternary graph showing a simplified cell wall composition based on their content of pectin, cellulose and Hyp-rich glycoprotein with extremes ranging from the pollen tube tip (~100% pectin), secondary cell walls (nominally 100% cellulose) to Chlamydomonas (~100% Hyp-rich glycoprotein) with the primary cell wall of higher plants and the Charophycean alga Coleochaete representing intermediate values.

Anton Heyn’s great insight defined the essential role of cell wall plasticity in cell extension (Heyn, 1940). The novel idea of the cell wall as a true plastic reflected the plastics revolution and zeitgeist of the 1930s. Curiously the simple extrapolation from plastics to plasticiser has been ignored due to the lack of candidates and a general consensus requiring the cleavage of load-bearing bonds although these remain unidentified. Not surprisingly the molecular basis of plasticity has remained speculative with many competing hypotheses including: Insertion of pectin polygalacturonate as a chelator of Ca^2+^ crosslinks (Proseus & Boyer, 2006; Hepler *et al*., 2013) combined with fluctuations in apical stiffness ascribed to pectin demethylesterification (Zerzour *et al*., 2009) (Bidhendi & Geitmann, 2016). An alternative “acid growth hypothesis” (Kutschera, 1994) formulated almost fifty years ago (Rayle & Cleland, 1970) involves proton secretion and concomitant cleavage of putative acid labile polysaccharide crosslinks similar to the expansin hypothesis (McQueen-Mason & Cosgrove, 1994). Auxin-induced proton secretion is a major tenet of the acid growth hypothesis but it also increases cytosolic Ca^2+^ (Vanneste & Friml, 2013); this is consistent with exocytosis of AGP wall plasticizers (Schopfer, 1990; Kutschera & Niklas, 2007) during pollen tube growth. Microscopically the pollen tube tip wall appears as a single pectin layer ~100 nm in width equivalent to >100 monomolecular layers of highly methylesterified pectin that are interspersed with AGPs. The Yariv reagent shows AGPs concentrated at the pollen tube tip (Mollet *et al*., 2002) (Jauh & Lord 1996) but also appearing as rings along the pollen tube (Li *et al*., 1992). Classical AGPs are ideal pectic plasticizers. By analogy with synthetic plasticisers that disrupt the orderly alignment of linear polymers, the intercalation of the small bead-like Hyp-arabinogalactan glycosubstituents (Hyp-AGs) (Lamport *et al*., 2014) likely disrupts linear pectin alignment. Indeed, direct experimental evidence involves the **Yariv reagent that inhibits tip growth with a concomitant rapid accumulation of periplasmic AGPs (Mollet *et al*., 2002); arguably a decreased level of AGPs *in muro* is causally connected with decreased wall plasticity**. We also infer that small arabinogalactan peptides may also ***increase wall plasticity*** based on their dramatic upregulation during auxin induced root cell elongation (Pacheco-Villalobos *et al*., 2016). Compared with the much larger classical AGPs (>100 kDa), the higher diffusibility of small (~20 kDa) AG-peptides (Van den Bulck *et al*., 2005) presumably enables them to plasticize thicker walls than at the pollen tube tip.

Indirect experimental evidence also strongly correlates AGPs with pollen tube tip growth (Seifert & Roberts, 2007) and also with more general root epidermal cell expansion (Ding & Zhu, 1997). Thus, double null mutants of pollen-specific *agp*6 and *agp*11 yielded pollen grains highly defective in germination and growth (Coimbra *et al*., 2009). Anecdotal evidence also correlates the friability of cell suspension cultures with enhanced AGP secretion.

Interestingly, when the Yariv reagent inhibits normal growth and tip extension ceases the pollen tube does not rupture, presumably because further additions such as callose thicken the tip wall (Jauh & Lord, 1996).

Figure 2 combines wall biomechanics and biochemistry by integrating AGPs and Hechtian adhesion sites as essential components of a Hechtian oscillator. These components physically connect the wall with stretch-sensitive components of the plasma membrane that control the Ca^2+^influx essential for coordinating the exocytosis of wall precursors. Thus, ***the wall regulates its own growth by Hechtian feedback***.

## Pollen tube guidance from stigma to embryo sac

A plethora of guidance cues that may direct pollen tube growth include: K^+^ Cl^−^ Ca^2+^ (Hepler *et al*., 2006), glycoproteins (Sommer-Knudsen *et al*., 1998), reactive oxygen species (ROS) (Foreman *et al*., 2003), nitric oxide (Prado *et al*., 2016), peptides (Qu *et al*., 2015) and complex signalling networks that remain to be elucidated (Leydon *et al*., 2015). However other guidance cues can now be considered from the perspective of the female reproductive tract where AGP-rich regions visualised by anti-AGP monoclonals (Coimbra & Salema, 1997; Coimbra & Duarte, 2003) coincide with the pathway traversed from stigma to the egg cell. Indeed, the remarkable coincidence of AGPs and Ca^2+^ throughout the female reproductive tract (Coimbra & Duarte, 2003) is hardly fortuitous. Faced with competing hypotheses Occam’s Razor suggests a simple Ca^2+^ guidance cue: (Fig. 6) Pollen tubes grown in vitro acidify their growth medium (Feijo *et al*., 1995); therefore in vivo they presumably dissociate AGP-Ca^2+^ of the transmitting tissue thus enabling pollen tubes to blaze a Ca^2+^ trail to the ovule. A dual source of cytosolic Ca^2+^seems likely: Firstly, at the tip derived from its periplasmic AGP-Ca^2+^ capacitor that involves recycling or “reflux” of cytosolic Ca^2+^. Secondly, Ca^2+^ released from surrounding tissues by the marked proton efflux at the pollen tube shank rather than at the tip (Hepler *et al*., 2006). Due to the cooperative effect of multiple pollen tubes (Heslop-Harrison *et al*., 1985) enhanced H^+^ efflux and release of Ca^2+^ from the locally abundant AGP-Ca^2+^ could contribute to reproductive success by ensuring fertilisation of multiple ovules. Evidence for the co-localisation of AGPs and Ca^2+^ at each stage of the pollen tube pathway follows:

**Fig. 6.**
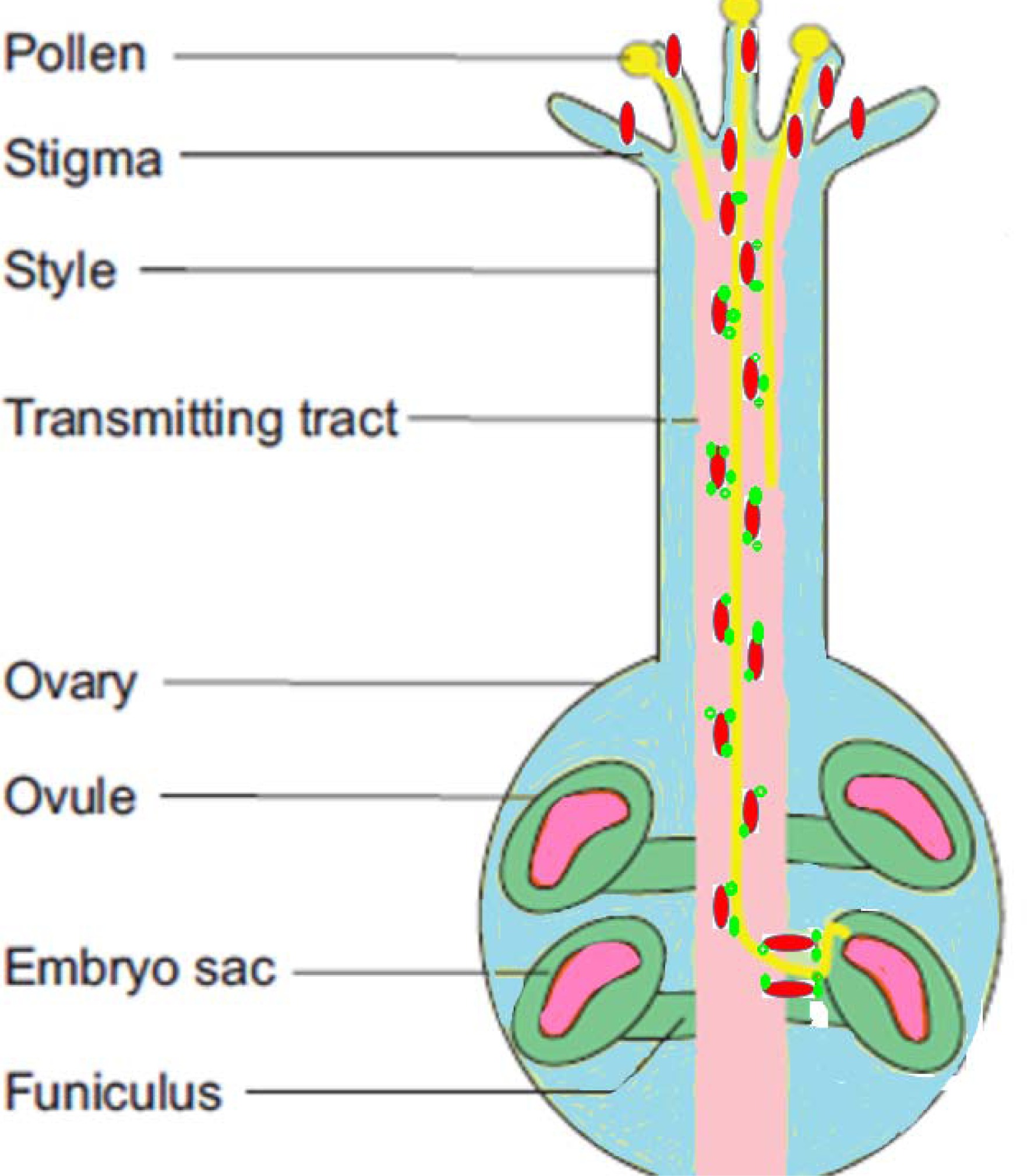
Pollen tube pathway to the embryo sac. Redrawn from (Qu *et al*., 2015) to show pollen tubes (yellow) growing through transmitting tissue lined with AGP-Ca^2+^ (red ellipsoids with green Ca^2+^) Pollen tubes signal to the transmitting tissue to supplement their endogenous Ca^2+^ levels by analogy with marathon runners who signal for a water bottle at stations along the track.

### Stigmatic tissue

Calcium antimonate histochemical detection of abundant Ca^2+^ in stigmatic tissue of several species (Ge *et al*., 2007) parallels AGPs detected by the β-D-Yariv reagent and also by monoclonal antibodies JIM8 and JIM13. Significantly, in apple blossom, stigmatic receptivity was acquired concomitantly with the secretion of AGPs (Losada & Herrero, 2012).

### Stylar tissue

Classic work (Mascarenhas & Machlis, 1962) showing Ca^2+^-dependent pollen tube growth and an elevated calcium content of tissues from stigma and transmitting tract of the style to the ovule has been amply confirmed (Knox *et al*., 1976), including Ca^2+^ chemotropism (Malho & Trewavas, 1996). As the distribution of AGPs and Ca^2+^ in the style coincide, the role of AGPs as a primary source of cytosolic Ca^2+^ seems likely.

### Ovule micropyle: synergids and filiform apparatus

On its final path towards the egg cell the pollen tube makes a sharp turn from the transmitting tissue toward the micropyle (Coimbra & Duarte, 2003) (Fig. 6) guided by signals from the synergid cells and filiform apparatus both strongly expressing AGPs (Coimbra & Salema, 1997) and Ca^2+^ (Chaubal & Reger, 1990) with highest levels of Ca^2+^ in the synergid filiform apparatus (Ge *et al*., 2007).

Tip growth is a fine balance between cell wall thickening at high levels of Ca^2+^ (Picton & Steer, 1983) and tip bursting that occurs in low Ca^2+^ (Hill *et al*., 2012) and increased auxin levels (Zerzour *et al*., 2009). However tip bursting at low oxygen levels is of physiological significance in the hypoxic ovary where failure of tip wall integrity (Linskens and Schrauwen 1966) releases gametes that fertilise the egg cell.

Significantly AGP expression is absent from the rudimentary ovules of male flowers of *Actinidia deliciosa* (Coimbra & Duarte, 2003) while laser ablation showed that only a single synergid cell in the isolated embryo sac of *Torenia fornieri* was necessary and sufficient for pollen tube attraction to the female gametophyte (Higashiyama *et al*., 2001). However, additional signals likely involve an AGP sidechain fragment designated AMOR or “Activation Molecule for Response” by the pollen tube to LURE guidance peptides from the synergids (Okuda *et al*., 2009). However, AMOR was identified as methyl-O-glucuronosyl-β-D-galactose (Mizukami *et al*., 2016) that is a key component of the AGP Ca^2+^-binding motif and thus suggests its possible role as a Ca^2+^ carrier. This complements the invariable coincidence of AGP and Ca^2+^. Clearly Occam is a guide not a guarantor!

## Future research pathways guided by Occam and Darwin

In 1799 Napoleon’s troops entered the Egyptian village of Rosetta (Rashid) and discovered an ancient basalt slab with a trilingual inscription in praise of King Ptolemy V (205-180 BC) that finally enabled Champollion to decipher Egyptian hieroglyphics in 1822. Rosetta is a metaphor for AGPs whose primary function remained unknown for fifty years until their structural hieroglyphics (Fig. 3) were deciphered to reveal the molecular function as an AGP-Ca^2+^ capacitor at the cell surface. This leads to a unified role for tip-localised AGPs and tip-focussed Ca^2+^ in cell extension as proposed here. However, viewed as a true plastic the control of wall plasticity at the molecular level in particular appears to be an intractable problem with widely differing views and assumptions. Nevertheless, the properties of AGPs combined with Hechtian adhesion offer a solution based on a Hechtian oscillator that generates cytosolic Ca^2+^ oscillations. This role for classical AGPs depends on their precise cell surface location and their unique chemistry: an exquisitely designed glycomotif with paired glucuronic acid residues that bind Ca^2+^ but released on demand. Such elegance would have pleased Paley but puzzled Darwin. Many questions remain. They include evolutionary origins (Verret *et al*., 2010). A vital clue lies embedded in the chalk cliffs of the South Downs National Park built over two hundred million years of fossilised phytoplankton; the calcified cell walls of coccolithophores, typically *Emiliania huxleyi*, may be the evolutionary precursor to dynamic Ca^2+^ storage at the cell surface of higher plants possibly with even wider implications of the Hechtian oscillator as an osmosensor (Haswell and Versluis 2015) or a gravisensor (Schnabl 2002) both in search of their identity as nanoscale molecular devices.

Finally, the reason for the wide range in classical AGP structure that includes the complex glycosylation of its protein core. Clearly the tiny minority of AGPs characterised biochemically only hints at their true diversity and versatility emphasised by recent bioinformatics that show higher plants have heavily invested in AGPs (Ma et al. 2017). Thus diverse stimuli might generate specific Ca^2+^ signatures (Rudd and Franklin-Tong 2001) based on the distribution, size and composition of AGP glycomodules. Because AGP glycosylation is only indirectly coded by the genome, precise structural oligosaccharide details cannot be predicted by bioinformatics. Hence the technical problem of rapid polysaccharide/oligosaccharide structural analysis and the determination of Ca^2+^ binding constants. Future *ab initio* computer simulations (cf. Fig. 3.) will enable the design of novel AGP glycomodules with properties optimised for a given environment in the perpetual quest for the Holy Grail of plant growth.

## Acknowledgements.

Professor Marcia Kieliszewski, Ohio University, for advice and encouragement. The University of Sussex, School of life sciences, for past laboratory facilities. Pembroke College, University of Cambridge, academic home of DTAL (1955-1961) and also William Turner (first English Herbal) and Nehemiah Grew (Father of plant anatomy).

DTAL is the corresponding author.

LT’s structural work made this paper possible and contributed to paper preparation.

MAH contributed to earlier structural work and paper preparation.

MJK has contributed many years work to this project and has provided invaluable advice.

